# Visual exposure enhances stimulus encoding and persistence in primary cortex

**DOI:** 10.1101/502328

**Authors:** Andreea Lazar, Christopher Lewis, Pascal Fries, Wolf Singer, Danko Nikolic

## Abstract

The brain adapts to the sensory environment. For example, simple sensory exposure can modify the response properties of early sensory neurons. How these changes affect the overall encoding and maintenance of stimulus information across neuronal populations remains unclear. We perform parallel recordings in the primary visual cortex of anesthetized cats and find that brief, repetitive exposure to structured visual stimuli enhances stimulus encoding by decreasing the selectivity and increasing the range of the neuronal responses that persist after stimulus presentation. Low-dimensional projection methods and simple classifiers demonstrate that visual exposure increases the segregation of persistent neuronal population responses into stimulus-specific clusters. These observed refinements preserve the representational details required for stimulus reconstruction and are detectable in post-exposure spontaneous activity. Assuming response facilitation and recurrent network interactions as the core mechanisms underlying stimulus persistence, we show that the exposure-driven segregation of stimulus responses can arise through strictly local plasticity mechanisms, also in the absence of firing rate changes. Our findings provide evidence for the existence of an automatic, unguided optimization process that enhances the encoding power of neuronal populations in early visual cortex, thus potentially benefiting simple readouts at higher stages of visual processing.

## Introduction

A key property of cortical circuits is their capacity to reorganize structurally and functionally with experience (1–3). In primary visual cortex, adaptive reorganization is well documented during development (4–7) and growing evidence indicates that sensory responses continue to adapt in adulthood (8–13). The continual refinement of sensory neurons based on the statistics of the sensory environment is at odds with the traditional view of the primary visual cortex as a collection of static filters or feature detectors, passively converting sensory input into a sparse code for further feed-forward processing across the visual hierarchy (14). In fact, considerable evidence suggests that primary visual cortex does not statically encode the environment but has rich spatial and temporal dynamics. For example, sensory-evoked activity propagates through the local network in wave-like patterns (15–17), displays a high-degree of temporal structure (18) and can persist long after the cessation of stimulation (19–22). These rich dynamic properties exhibited by early visual neurons suggest an active involvement of primary visual cortex populations in the coordinated representation of visual stimuli. Most strikingly, repetitive visual exposure can alter the strength and selectivity of neuronal responses in the primary visual cortex, leaving a lasting mark on post-exposure activity in both awake and anesthetized animals (23, 24). Yet, it remains unclear how such changes affect the joint encoding of stimuli across neuronal populations and ultimately the information transmitted to downstream areas.

Given that primary neurons adapt their responses as a function of repeated exposure, one compelling hypothesis is that exposure-driven changes are coordinated across neuronal populations to collectively improve the representation and maintenance of recently experienced stimuli. Here, we test this hypothesis by investigating the impact of visual exposure on the persistent population response of neurons in cat area 17 to brief, structured stimulation. We employ a large set of abstract stimuli (letters of the Latin alphabet and Arabic numerals) which provide a rich variety of spatial conjunctions across low level features and are well suited to capture aspects of distributed coding. We find five main signatures of functional reorganization. First, visual exposure optimizes stimulus maintenance in primary visual cortex by increasing the magnitude and decreasing the variability of neuronal responses that persist after stimulus offset. Second, these changes are associated with neural recruitment, a broadening of the dynamic range neurons employ to respond to stimuli, and an enhancement of stimulus-specific tiling of neuronal responses. Third, refinement of individual responses results in increased stimulus encoding at the population level, i.e. a simple hypothetical downstream decoder increases its accuracy in identifying recent stimuli from brief snippets of population activity. Fourth, the exposure-driven enhancements in stimulus persistence maintain the representational structure of stimuli resulting in improved stimulus reconstruction. Fifth, exposure strengthens patterns in postexposure spontaneous activity. Finally, modeling demonstrates that exposure-driven enhancements in stimulus persistence can arise from recurrent network interactions via local, unsupervised plasticity-mechanisms.

## Results

We used silicon multi-electrode arrays to record the simultaneous activity of neuronal populations in area 17 of five lightly anesthetized adult cats (*felis catus*; mean age 2.7 years; range 1-5 years; two females).We applied standard spike-sorting techniques to isolate the action potentials of 112 single-unit and 331 multi-unit clusters (Materials and Methods). The receptive fields (RFs) of the recorded units (27-52 units per session, 11 recording sessions with independent electrode positions, 443 units in total) were located nearby in visual space and were jointly stimulated by a single luminance stimulus, flashed for 100 milliseconds over a black background (example trials in Figure 1A). Short stimulus presentations at high contrast can produce strong persistent responses in the primary visual cortex (20, 21). In our data, the flashed stimuli evoked a biphasic population response, composed of a transient, low-latency component (≈ 50 ms) and a prolonged, persistent component (example trials and average firing rates across all 443 units from 5 cats in Figure 1A). In total, 296 out of 443 units (66.8%) fired above the baseline for the entire duration of the trial (comparison between the last 100 ms of the trial and the pre-stimulus baseline, one-tail t-test, p<0.05).

**Fig. 1.**
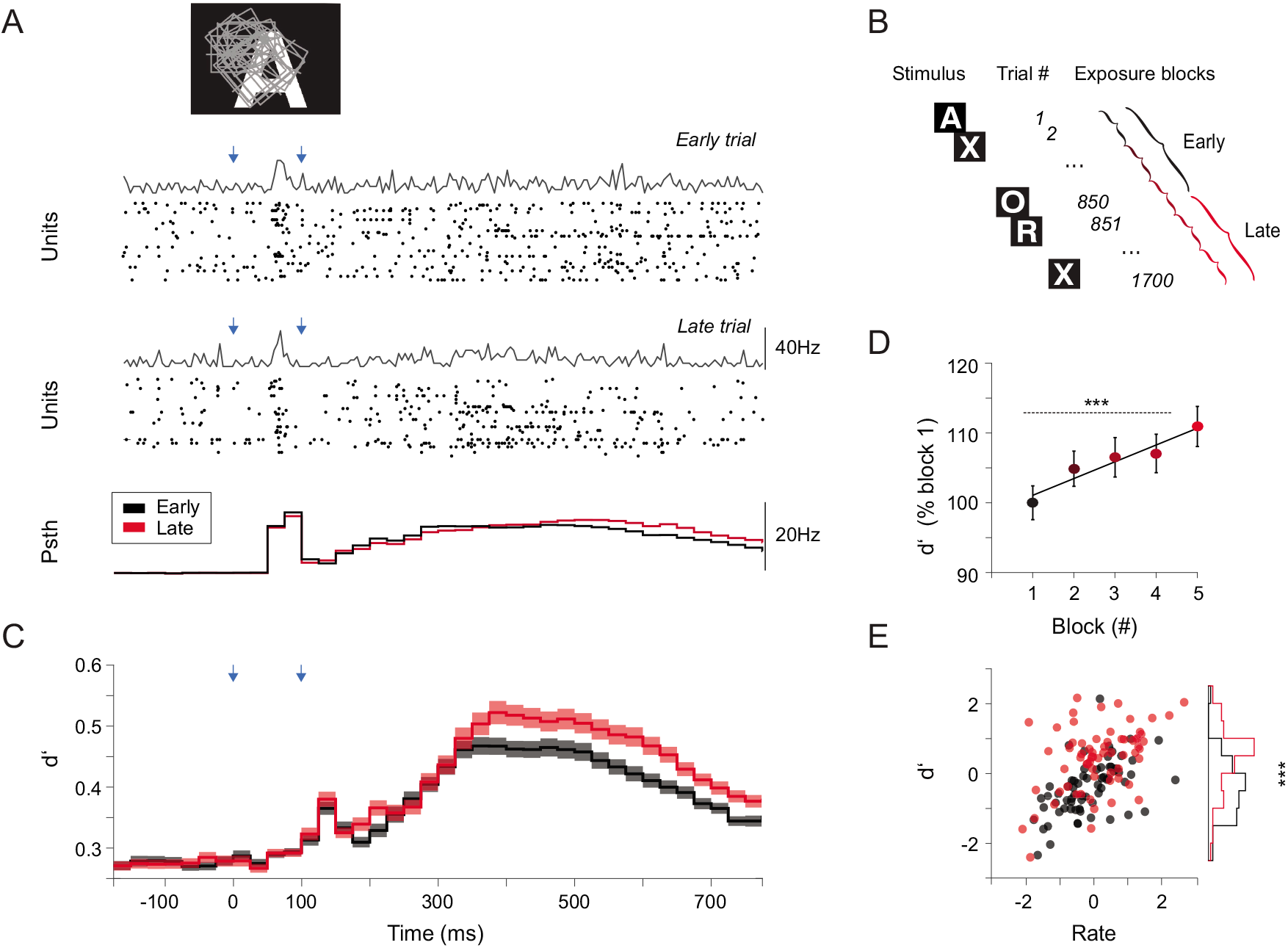
Visual exposure protocol and simultaneous recordings of neuronal population activity from cat area 17. A) Cluster of receptive fields of simultaneously recorded multi-units (rectangles) relative to the location of a visual stimulus. Population activity in example trials from one session. After a brief stimulus presentation (100 ms, on and off timing marked in blue arrows), neuronal responses display a short transient followed by a persistent reverberatory component. Bottom peristimulus time histogram shows mean population firing rates across 443 recorded units in early (black) and late trials (red). B) Exposure protocol: in each session 34 visual stimuli were presented in random order (1700 trials in total). Sessions were split in either two or five consecutive blocks of trials and analyzed separately. C) Average stimulus discriminability (d’) over the course of the trial (d, calculated per unit; 443 units; 34 stimuli; shaded area indicates the standard error from the mean). The effect of visual exposure on stimulus discriminability is significant for the persistent (300-600 ms), but not the transient(0-300 ms) part of the evoked response. D)The increase in d’ is gradual and does not saturate for the exposure interval considered here (5 blocks; interval 300-600 ms). E) Similar firing rate levels result in higher discriminability in late trials (red) compared to early trials (black) in a session. In all subplots *** stands for p < 0.001.

The anesthetic protocol used here, consisting of intravenous suffentanil supplemented by minimal concentrations of isoflurane, was intended to model stable cortical dynamics, absent of strong fluctuations between ‘up’ and ‘down’ states. The stability and quality of the recordings was quantified using the power spectrum of the local field potential (LFP), and comparing the shapes of early and late spikes, neither of which exhibited any systematic change over the exposure interval (Figure S1).

### Visual exposure enhances stimulus persistence

How does brief visual exposure to structured stimuli affect the population response of primary neurons? To address this question, we presented a large set of alphanumeric stimuli (34 upper-case letters and digits) in random order (1700 trials in total, 50 trials per stimulus) and compared stimulus responses across either two or five consecutive trial-blocks (stimulus order within each block was random, consecutive trials corresponded to different stimuli, schematic in Figure 1B).

We measured stimulus discriminability, also known as Cohen’s *d*’ (25), by calculating, for each individual unit and each 50 ms time window within the trial, the spread of the mean responses to different stimuli relative to the standard deviations of those responses across trials (definition of *d*’ for 34 stimuli in Materials and Methods). We found that visual exposure led to a substantial increase in average *d*’ across units (2 blocks, 8.38% increase for the interval 100-800ms, paired t-test, p = 6.4e-13,t = −7.43, df = 442; 10.86% increase for the interval 300-600 ms, paired t-test p = 2.8e-12, t = −7.19, df = 442; profile of average *d*’ along the trial in Figure 1C). The increase in *d*’ with visual exposure was gradual and did not reach a saturation point (5 blocks, 10.9% increase between block 1 and 5, paired t-test p = 1.3e-07, the black line indicates the linear fit, y = 2.4x + 98.66, linear trend was significant p = 0.009; Figure 1D), suggesting that further improvements may be possible with further exposure.

An improvement in stimulus discriminability is likely to be associated with an increase in neuronal response amplitude or a decrease in response variability. We found that visual exposure resulted in an increase in the amplitude of neuronal responses that persisted after stimulus presentations (6.2% increase between early and late trials for the 300-600ms interval; paired t-test, p = 2e-08, t = −5.71, df = 442; see Figures 1A and S2). Interestingly, similar firing rate levels resulted in higher *d*’ values for late trials in a session, suggesting that modulations in firing rates alone cannot explain the observed improvements in stimulus discriminability (Figure 1E). Moreover, the same conclusion was reinforced by the observation that, in two animals, the mean population firing rate was unchanged by stimulus exposure, in spite of substantial improvements in stimulus encoding (cat 3 and 5, see related stimulus decoding performance in Figure S5). It is known that firing variability is reduced by stimulus onset (26). Here we found that visual exposure further reduced variability throughout the trial (3.48% decrease between early and late trials for the interval 300-600 ms; paired t-test, p = 8e-05,t = 3.98, df = 442, Figure S2). Laminar analysis revealed that exposure driven changes in response amplitude, variability and *d*’ were significant for all compartments (Figure S3).

### Exposure increases the dynamic range and stimulus–clustering of neuronal responses

Given the common assumption that higher response selectivity corresponds to more stimulus information, we considered the possibility that the observed exposure-driven enhancement in stimulus discriminability may be associated with an increase in response sparseness or selectivity. We assessed both the population sparseness for each stimulus, and the stimulus selectivity of each unit, separately for early and late trials in each session (response period 300-600 ms, see Material and Methods). Sparseness was estimated as one minus the fraction of simultaneously recorded neurons that responded to each stimulus, and, conversely, the selectivity of each unit was estimated as one minus the proportion of stimuli it responded to. We found that both measures decreased with visual exposure (sparseness, paired t-test across stimuli and sessions p= 6.8e-04, t = 3.4, df = 373; values z-scored per session; Figure 2A; selectivity, paired t-test, p = 0.006,t = −2.7, df = 442; not shown). When units were sorted based on their change in firing rate amplitude and grouped into quartiles, we found that the units that strongly decreased their firing rates with exposure, showed increased selectivity, but decreased d’ (compare Figure 2B and C). Conversely, the units that increased their firing rates with exposure became less selective and increased their d’ values. Interestingly, we found that exposure recruited more units to stimuli (reduced sparseness) and that recruited units increased the dynamic range of their firing rate responses (the difference between the strongest and the weakest response across stimuli, see Material and Methods, Figure 2D). The increased dynamic range was highly significant across units (paired t-test, p = 7.5e-17, t = −8.6, df = 442).

**Fig. 2.**
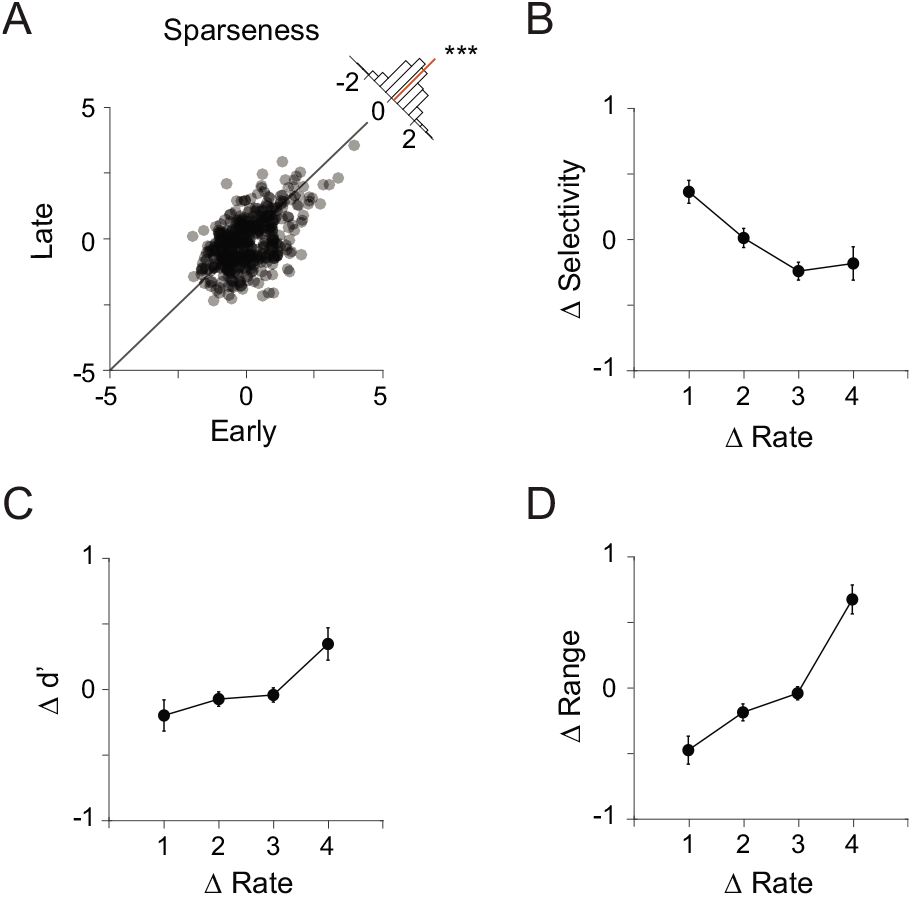
Exposure-driven refinements in stimulus-encoding A) Response sparseness decreases significantly with visual exposure, suggesting neural recruitment. Individual markers correspond to individual stimulus conditions (11 sessions, 34 stimuli). B) Change in response selectivity as a function of change in firing rate with exposure. Units are sorted by amplitude of rate change and grouped into quartiles (marked on x-axis). Values on y-axis are z-scored per session. C) Change in *d*’ as a function of change in firing rate with exposure. Units that increase their firing rates with visual exposure also increase their *d*’ values, but are accompanied by a loss in selectivity (compare to B). D) Change in response range as a function of change in firing rate with exposure. Positive gains in response range are associated with an increase in firing rates. All measures are calculated over the 300-600 ms time interval in the trial. Error bars in B, C and D indicate standard errors from the mean.

Reducing the high-dimensional population response via principal component analysis (PCA), we found that exposure increased the segregation of responses, revealing stimulus-specific clusters in low-dimensional projections (examples in Figure 3A and B; multiple sessions in Figure S4). Better segregation was quantified as reduced cluster radius and increased inter-cluster distance. The cluster radius, the mean Euclidean distance in the first two principal components of all cluster-points (stimulus trials) to the cluster center (average response), decreased significantly with exposure (10.4% decrease, paired t-test p= 1.25e-15,t = 8.36, df = 373; Figure 3C). In addition, the cluster distance, the mean distance between the cluster center and all other centers, increased with exposure (15.3% increase, paired t-test p= 4.7e-25, t= −11.13, df = 373; Figure 3D). Overall, the clustering index, calculated per session as one minus the mean ratio between the cluster radius and the cluster distance, increased with exposure (paired t-test p= 0.0058,t = −3.48, df = 10; Figure 3E).

**Fig. 3.**
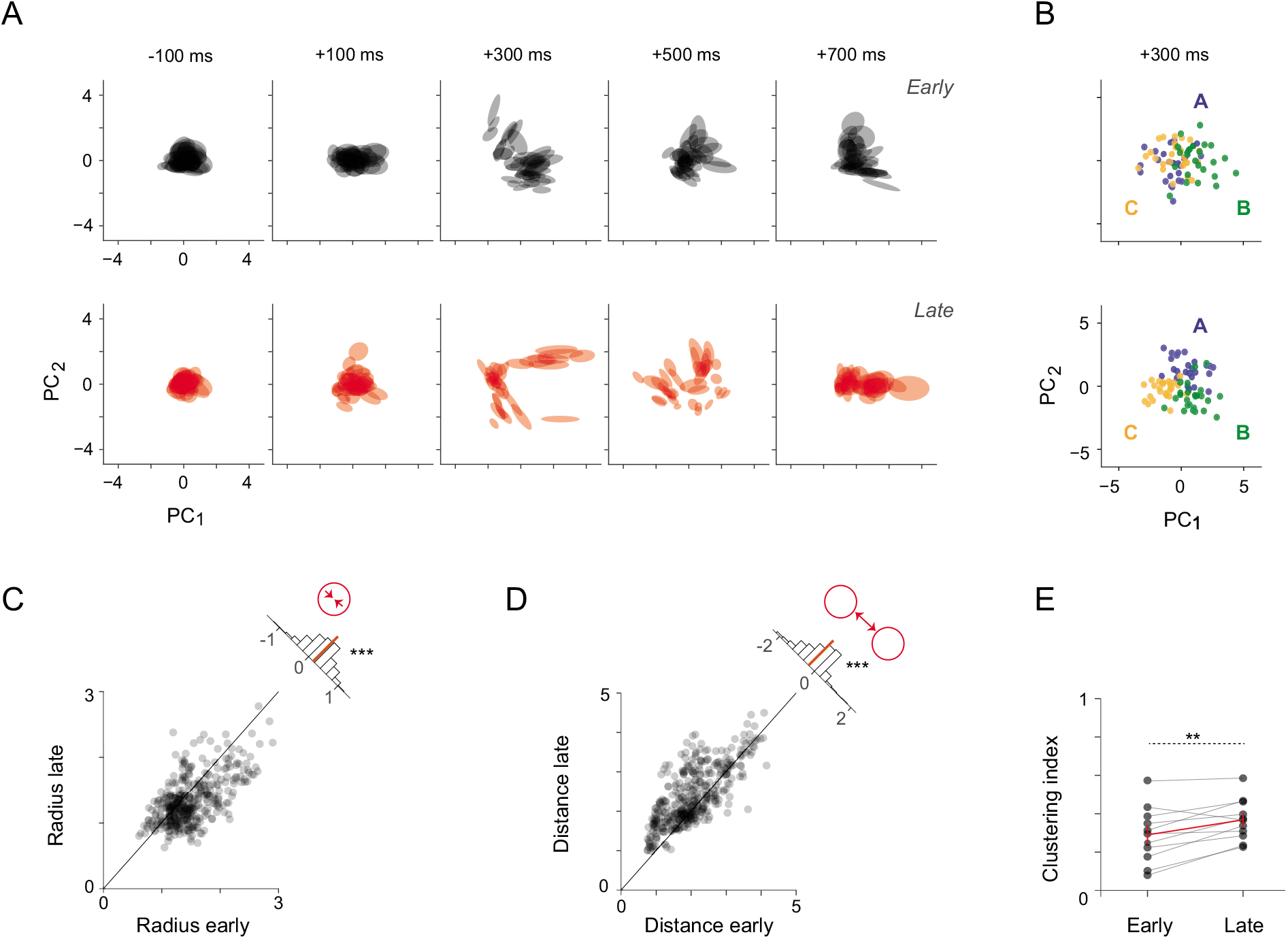
Exposure increases stimulus-specific clustering and segregation of the population responses. A) Evolution of population responses to 34 visual stimuli over the course of the trial (example session, 50 ms spike-count vectors). B) Early and late population responses (50 ms spike-count vectors) to three stimuli (letters A, B and C) in the space defined by the first two principal components. Each marker represents a single trial. For each cluster, an ellipse circumscribes the data points within one standard deviation from the mean. The stimulus-specific clusters segregate ≈ 300 ms after stimulus onset. The segregation is more pronounced for late trials (red) compared to early trials (black) in a session. C) Scatter of cluster radius values in early and late trials, for all stimulus clusters in all sessions (50 ms spike-counts, 300 ms after stimulus offset). The small inset histogram shows the distribution of differences in cluster radius between early and late trials, the mean of the distribution which is significantly different from 0 is marked in red. D) Scatter of mean distances from each cluster center to all others, in all sessions (same time window as above). The inset histogram shows the distribution of differences in cluster distance between early and late trails. E) The clustering index increases significantly across recording sessions with visual exposure.

In sum, these results suggest an exposure-driven refinement of stimulus encoding. This refinement does not occur through increased selectivity of units, or population sparseness, but rather through the recruitment of more responsive units into the population response, an expansion of the dynamic range of units, and enhanced stimulus-specific clustering of population responses.

### Exposure enhances readout performance

We next sought to investigate the extent to which exposure-driven changes in neuronal responses affect the capacity of a hypothetical downstream decoder to identify visual stimuli based the primary visual cortex output.

We trained independent Bayesian classifiers to perform time-resolved decoding of stimulus identity based on the population activity vector across the trial, i.e. the spike-count in each time bin (instantaneous decoders, schematic in Figure 4A). Visual exposure led to increased classification performance (time course in Figure 4B, chance level = 2.94% for 34 stimuli, 50 ms time bins, 100-fold validation procedure, see Materials and Methods for details). The magnitude of the increase was substantial given the modest changes in firing rate and variability observed for individual units. Peak accuracy across sessions ranged from 8 to 49.5% correct for early trials and 16.5 to 59.6% correct for late trials, and increased significantly in every animal (average increase 27.7%, range 13-59.6% increase, t-test, all p-values<0.001). To quantify stimulus persistence, we analyzed the time course of classification performance within the trial by calculating the area under the curve (AUC, 100-800 ms). We found that visual exposure led to a strong and significant increase in performance AUC in every animal (average increase 33%, range 14-64%, t-test, all p-values<0.001, individual performance profiles in Figure S5). Given the large number of stimuli in our set, individual stimuli were typically repeated only 50 times. Previous studies have indicated that remarkably little stimulus-exposure is required to modify the response properties of neurons in the primary visual cortex (24). However, to test the effect of more repetitions, we acquired a session of 5100 trials (150 trials per stimulus). In this control session, the performance AUC continued to increase past 50 repetitions per stimulus (Figure S6), suggesting that additional exposure continues to enhance stimulus encoding.

**Fig. 4.**
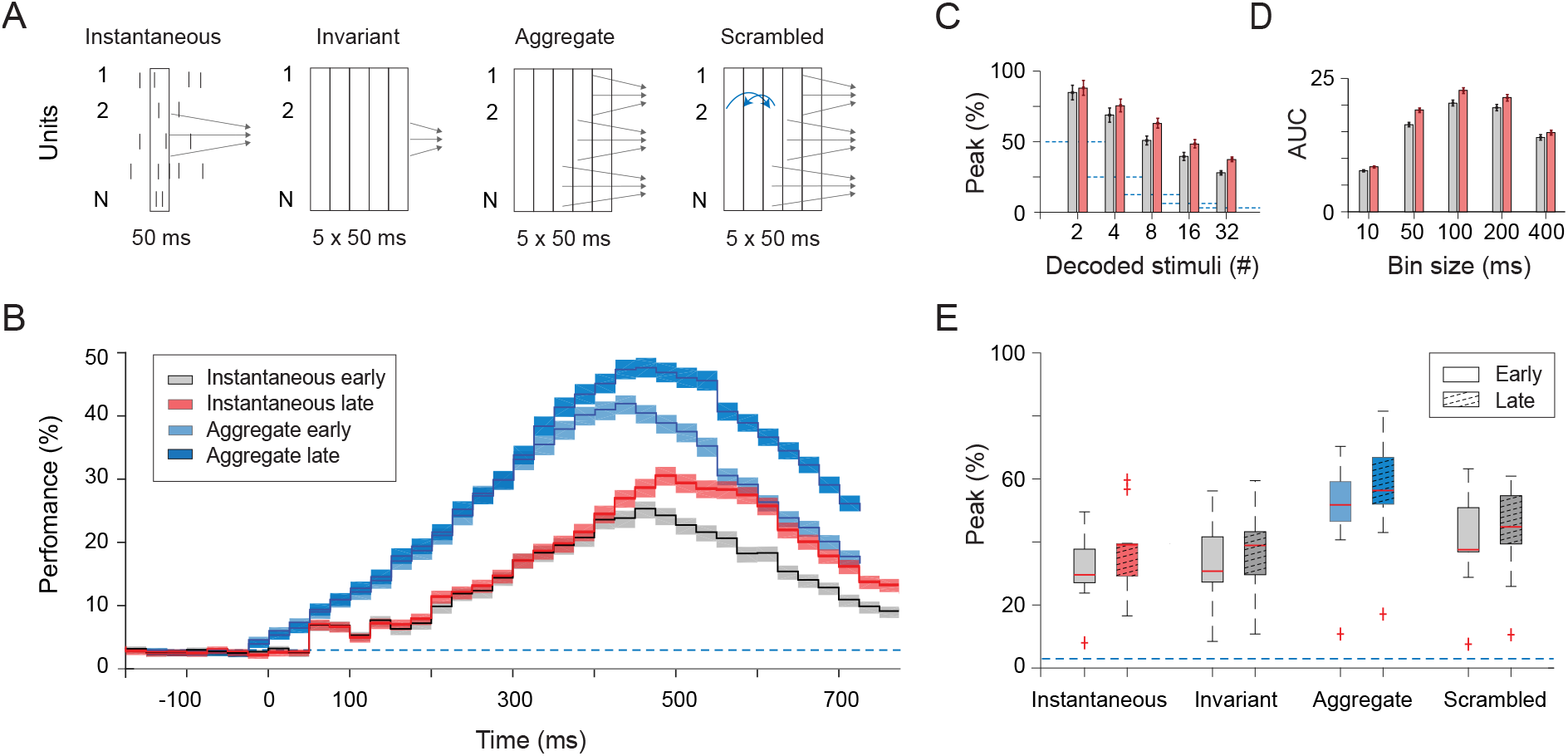
Exposure enhances stimulus decoding performance A) Types of Bayesian decoders: instantaneous (N-dimensional readout trained and tested on brief 50 ms spikecounts, independently for each time bin), invariant (*N*-dimensional readout trained on population spike-counts from five consecutive 50 ms time-bins; quantifies information that is consistent across trial time), aggregate (5 × *N* dimensional readout integrates information from five consecutive time-bins), scrambled (5 × *N* dimensional readout integrates across five time bins after temporal-scrambling). B) Stimulus classification performance over the course of the trial for instantaneous and aggregate decoders (chance level indicated by dotted line, individual sessions in Figure S5). C) Peak classification performance of instantaneous decoder as a function of task difficulty (number of stimuli being classified), relative to chance level (dotted lines). Differences between early and late trials are more pronounced for harder classification problems. D) Dependence of performance of instantaneous decoder, quantified by the area under the curve (AUC), on the integration bin size denoted on the x-axis, for early and late trials (upper panel). Performance is highest for intermediate integration intervals. E) Peak classification performance for the Bayesian decoders described schematically in A. All four types are significantly improved suggesting that both variant and invariant aspects of stimulus encoding are enhanced by visual exposure

As expected, the bin size used to integrate spike counts affected the difference in decoding performance between early and late trials (Figure 4B). Decoders that counted spikes over intermediate integration windows (50-200 ms) had high performance and showed significant improvements between early and late trials (performance AUC, t-test, p<0.05), while very short (10 ms) and very long (400 ms) integration windows resulted in lower performance and reduced improvement (AUC, t-test, p>0.05). Additionally, the effect of visual exposure on decoding performance varied with the task difficulty, i.e. the number of stimuli being decoded (Figure 4C). We found that exposure improved peak performance when classifying 8 or more stimuli (t-test, p<0.05), but not fewer (t-test, p>0.05). This is likely due to ceiling effects as peak performance scores for 2 class problems were beyond 90% for early trials in 4 out of 11 sessions.

The segregation of evoked responses into stimulus-specific clusters varied substantially over the course of the trial (Figure 3A) and peaked at different moments in time for different animals (Figure S5). We therefore examined how stimulus-specific information varied with trial-time by considering three additional decoding configurations (schematic in Figure 4A). First, we trained “time-invariant” decoders on activity vectors pooled across five consecutive 50 ms time-bins. Note that such decoders have five times more data points for the same N-dimensional space, compared to the instantaneous decoders. Second, we trained “aggregate” decoders on concatenated activity vectors corresponding to five consecutive temporal bins. Aggregate decoders map a five times larger dimensional space compared to the instantaneous decoders. Finally, we trained “scrambled” decoders on temporally scrambled data across five temporal bins. Scrambled decoders have the same number of points and space dimensions as the aggregate decoders but map an altered space where within-trial correlations between neurons have been disrupted through scrambling. Interestingly, all three decoders showed significant changes with visual exposure. The aggregate decoder performed significantly better than both the invariant and temporally scrambled decoders suggesting that information was contained not only in the instantaneous structure of the spike-count vector but also in its trajectory (in the sequence of state vectors during trial).

Finally, we considered the impact of visual exposure on the portion of trial-to-trial variability shared between units. Consistent with previous studies (27, 28), spike-count correlations (SCCs) were highest for pairs of units with similar stimulus preferences (positive signal correlations) and lowest for pairs of units with opposing stimulus preferences (negative signal correlations). Visual exposure reduced the strength of SCCs (21% decrease, paired t-test, p =1e-17), and the reduction was strongest for units with opposing preferences (78% decrease, two-tailed t-test, p = 1e-09, signal correlations<-0.1; 9% decrease, two-tailed t-test, p = 1e-06, signal correlations>0.1; Figure S7A). Ignoring SCCs can decrease decoding performance (Averbeck et al., 2006; Graf et al., 2011). We found that a support vector machine with quadratic features trained on trial-shuffled data and tested on original data, performed worse than a decoder trained on the original data with intact correlation structure (two-way ANOVA; shuffling led to a 10.42 % decrease, p = 0.0062 early trials and 13.02 % decrease, p = 3.7e-09 late trials; exposure led to a 17.96% increase, p = 1.2e-07 for original data and 14.54% increase, p = 0.0003 for shuffled data; Figure S7B). Shuffling reduced performance for both the early and late trials suggesting that while repeated exposure decreased the overall level of SCCs in the data, a portion of SCCs present in both early and late trials contributed positively to the population code. Indeed, the fact that SCCs decreased most for units with opposing stimulus preferences might reflect competition between stimulus-specific ensembles, such that correlations are stable between units of similar preference and reduced between units of opposing ensembles.

### Exposure enhances stimulus reconstruction

The alphanumeric stimuli are structurally more complex than oriented gratings but less complex than natural scenes. Such a large stimulus set of intermediate complexity is highly suitable for reconstruction, i.e. recreating the luminance pattern of stimuli from their evoked neuronal responses. While stimulus decoding techniques have been applied to many visual cortical areas, stimulus reconstruction has been attempted rarely and not, to our knowledge, in the context of visual exposure.

We performed stimulus reconstruction separately on early and late trials to quantify the impact that visual exposure had on encoding. To reconstruct each stimulus, we trained Bayesian decoders to predict the luminance of individual stimulus patches (576 decoders corresponding to 24×24 image patches) based on the population activity recorded after stimulus offset (schematic in Figure 5A; luminance values were between 0 and 1; independent train and test trials, 20-fold validation scheme; 11 sessions). The reconstructed stimuli were noisy on single trials (examples from one session, 50 ms spike-counts, 400 ms after stimulus offset; Figure 5B left) and became considerably more accurate when averaged across 10 test trials (Figure 5B right). Exposure increased stimulus reconstruction accuracy, calculated as one minus the difference in pixel luminance between the reconstructed image and the original image (scatter of 34 stimuli from 11 sessions; paired t-test, p = 1.13e-09, t = −6.24, df = 373; Figure 5C left). This improvement was significant also when calculated across sessions (paired t-test, p = 8.2e-04, t = −4.71, df = 10; Figure 5C right).These results suggest that the exposure-driven enhancements in classification performance are representational in nature and reflect improved encoding of stimulus content. The fact that the structure of multiple stimulus shapes (34 letters and digits) can be reconstructed from relatively small populations of recorded units (27-52 units per session) and improves with exposure speaks to the impressive encoding capacity and flexibility of the primary visual cortex.

**Fig. 5.**
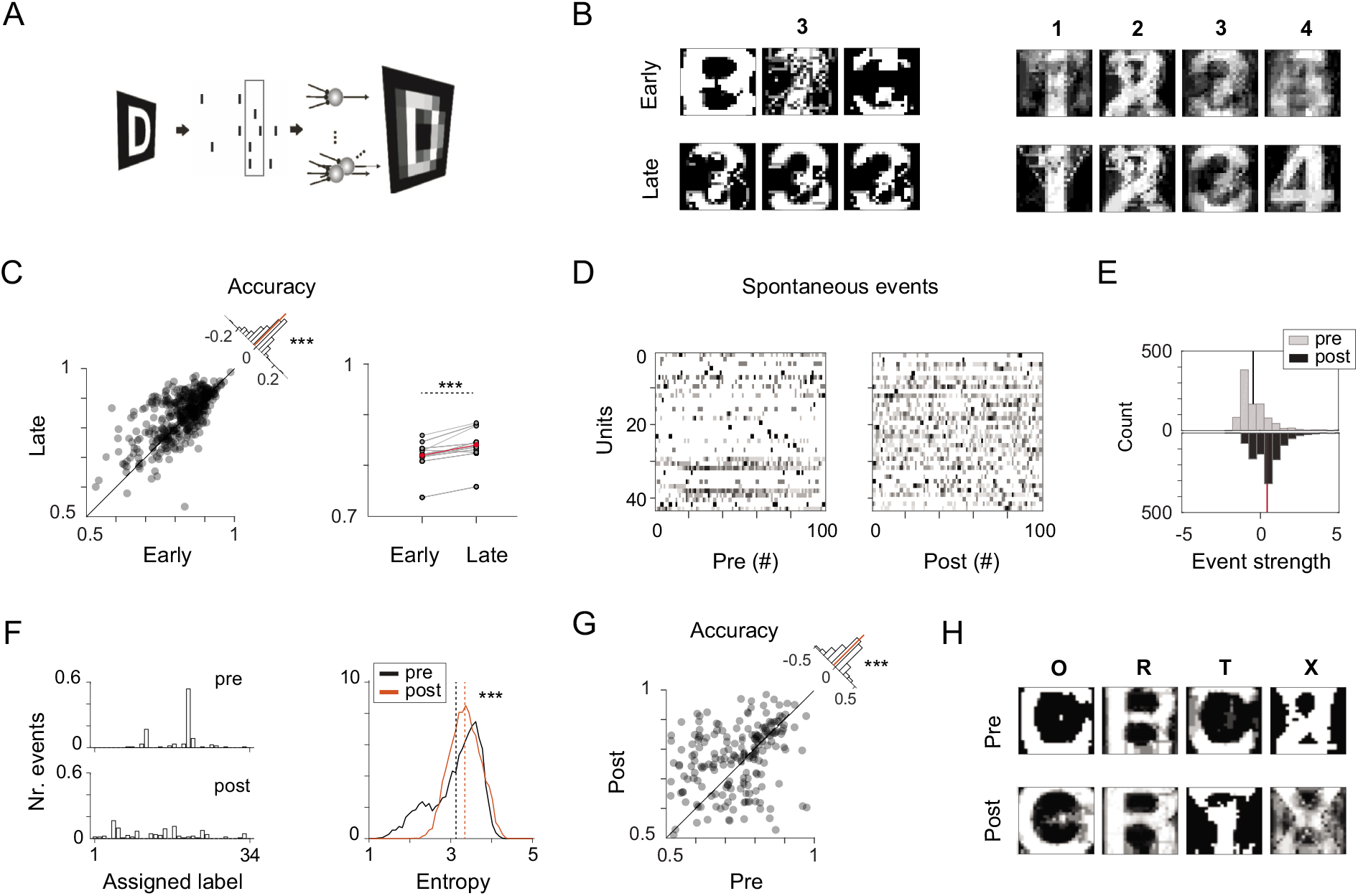
Improved stimulus reconstruction with visual exposure. A) Schematic representation of stimulus reconstruction technique: a 24×24 array of Bayesian classifiers are trained to predict pixel luminance from population spike-count vectors (50 ms) recorded 400 ms after stimulus offset. Test trials are omitted from the training set. B) Single trial examples for reconstruction of a stimulus (number “3”) and examples of average reconstructions (across 10 test trials, numbers “1” to “4”) for early and late trials in an example session. C) Scatter depicts the reconstruction accuracy of all stimuli from 11 sessions, early vs. late trials (left panel). Changes in reconstruction accuracy are significant (inset histogram, paired t-test, p<0.001; mean accuracy per session in right [panel, paired t-test, p<0.001). D) Example spontaneous events detected pre and post visual exposure in a session. E) Spontaneous events become stronger post-exposure (events detected pre and post exposure in 7 sessions). F) A Bayes classifier trained on evoked activity assigns stimulus labels to spontaneous events. The assigned labels are more uniformly distributed post-exposure; entropy is significantly higher post exposure (right panel; permutation test). G) Stimulus reconstruction accuracy based on spontaneous events improves post-exposure. H) Examples of single letter reconstructions based on spontaneous events occurring pre and post stimulus exposure (24×24 luminance patches; training on evoked data).

### Structured post-exposure spontaneous activity

Exposure-driven changes in evoked activity are often associated with accompanying changes in spontaneous activity. The stimulus reconstruction technique described above allowed us to probe for lasting representational changes in the structure of spontaneous neuronal activity. To this end, in 7 exposure sessions from 3 cats, we recorded and analysed spontaneous activity (20 blank trials) before and after visual stimulation.

We isolated spontaneously occurring strong-activation events, defined as periods when population activity exceeded mean activity by more than one standard deviation (50 ms spike-counts). In total, 961 events were detected during preexposure spontaneous activity and 980 during post-exposure (example events corresponding to pre- and post-exposure data from one session, Figure 5D). We found that the strength of spontaneous events increased significantly after visual exposure (event strength was pooled across sessions after being standardized for mean and variance; t-test p = 3.9e-89, t = −21,df= 1939; Figure 5E).

To quantify structural differences in the pre- and postexposure events, we assigned a stimulus ‘label’ to each event using decoders trained on evoked activity (example label assignments for pre- and post-exposure events from one session in Figure 5F). The assigned labels were more uniformly distributed across stimulus conditions post-exposure, as indicated by an increase in entropy (estimated based on 1000 bootstraps of 50 events, p = 1e-172, Figure 5F right). We next generated an image, using the same reconstruction technique as above, for each spontaneous event. Both the reconstruction and the identity decoders were trained on evoked activity from the entire session (no separation of early and late trials), as we wanted to assess exposure-driven changes in the structure of spontaneous activity, not changes in the reconstruction or decoding techniques. We pooled images across events based on the assigned class label to obtain mean stimulus reconstructions (reconstruction examples in Figure 5G). The accuracy of stimulus reconstruction improved following exposure (paired t-test, p = 5.3e-04,t = −3.52, df = 204; Figure 5H).Since the assignment of a class label and the reconstruction of an image for the same spontaneous event, reflect related content, it is not surprising that some degree of stimulus reconstruction is possible based on spontaneous activity. However, the improvement in post-exposure reconstruction accuracy suggests that the changes in spontaneous events are structured and correspond to the experienced visual content.

### Self-organized recurrent networks for stimulus persistence

Finally, we sought to demonstrate that a simple selforganized recurrent network, endowed with local plasticity mechanisms for learning and homeostasis, can qualitatively reproduce the exposure driven enhancements in stimulus encoding, while maintaining stable firing rate output.

The recurrent network model consisted of 80% excitatory and 20% inhibitory binary units (250 units total). For simplicity, the recurrent interactions were assumed to arise from a single pool of randomly connected neurons, not a multi-layered recurrent network. The connectivity matrix between excitatory units followed simple topological constraints, i.e. nearby excitatory neurons had increased probability for a connection (details in Materials and Methods). A subset of neurons received luminance input from a 6×6 array of stimulus image patches (schematic Figure 6A).

**Fig. 6.**
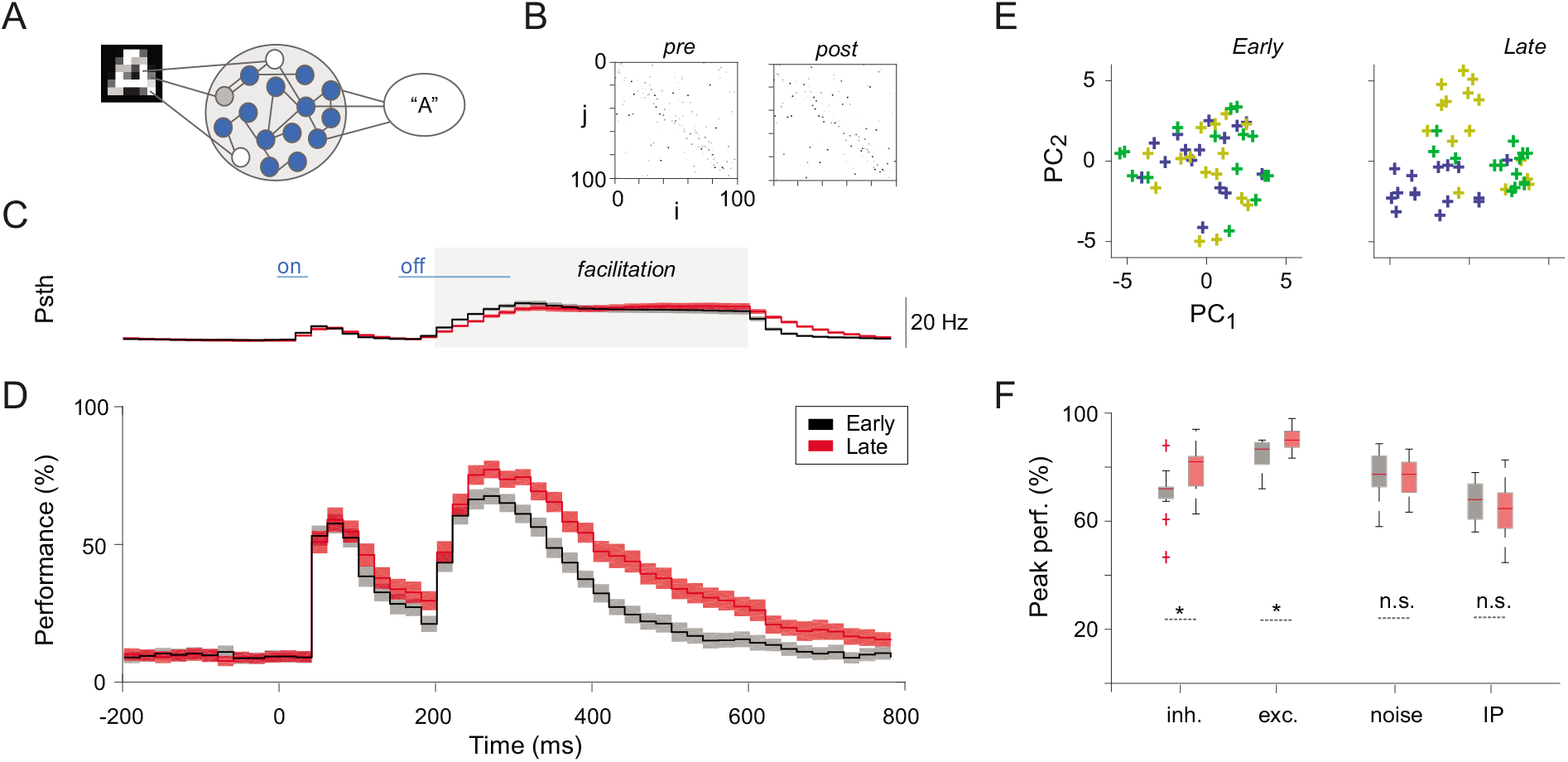
Self-organizing recurrent network (SORN). A) Recurrent neural network with subset of neurons receiving pixel luminance inputs. B) Network topology: initial connections are sparse and random with nearby connections more likely to occur. Example of connections between 100 excitatory units out of the 200 total, before and after learning in one simulation. Visual exposure resulted in self-organized changes in synaptic gains. C) Example of network excitatory response to a brief stimulus. Response facilitation, implemented here through a decrease in synaptic inhibition, is marked by gray area. D) Example of classification performance improvement with stimulus exposure via unsupervised local plasticity rules (shaded areas indicate standard errors from the mean across 10 simulation). Note that here the firing rates are similar in early and late trials, due to the homeostatic nature of intrinsic plasticity, thus the enhancement in encoding comes from a reorganization of internal network dynamics. E) Exposure results in an enhanced clustering of trials belonging to different stimulus conditions in a low-dimensional projection space. F) Facilitation can boost or impede unsupervised learning depending on implementation. Facilitation via a decrease in synaptic inhibition or an increase in synaptic excitation leads to a boost in classification performance. Facilitation via non-structured excitatory input (noise) or threshold modulation via intrinsic plasticity interferes with learning.

Recurrent neural networks naturally exhibit a memory of recent inputs, so information about brief stimuli can be retrieved with some delay from stimulus offset (29, 30). To match the empirical data, we strengthened the persistent recurrent responses after stimulus offset though response facilitation, which has been previously implicated in both stimulus persistence and learning (31, 32) (fixed interval for facilitation marked by shaded area in Figure 6C and D).

Visual exposure consisted of 50 brief presentations of 10 stimuli (alphabet letters A-J) in random order. The network self-organized through local, unsupervised synaptic plasticity, while homeostatic plasticity maintained neuronal firing rates at a fixed level (example of connectivity changes through plasticity in Figure 6B, details in Materials and Methods).

We considered how exposure-driven learning may interact with several implementations of response facilitation. Facilitation was implemented by changing the excitatory-inhibitory balance of incoming synaptic gains per neuron by either (1) lowering inhibition or (2) increasing excitation. Alternatively, the firing of excitatory units was increased by: (3) adding a random input drive and (4) changing neuronal excitability via intrinsic plasticity. The first two methods resulted in persistent firing rate responses after stimulus offset. Unsupervised learning during visual exposure led to improved stimulus encoding (data for an example run with lowered inhibition in Figure 6C,D and E). Similar to the empirical results, the enhancement in stimulus encoding could be captured in low-dimensional projections of the data (Figure 6E). The last two methods, variable input and changes in neuronal excitability, resulted in similar, persistent firing rate responses after stimulus offset. However, they did not lead to improved stimulus encoding through unsupervised learning, suggesting that the intrinsic network interactions, which were more severely disrupted by these two methods, played a critical role in the optimization process (Figure 6F).

## Discussion

We found that repeated exposure to briefly flashed visual shapes improves stimulus encoding in primary visual cortex. Visual exposure altered post-stimulus population activity in a manner that enhanced both the decoding of stimulus identity and the reconstruction of visual stimuli.

These improvements were associated with neuronal recruitment, an increase in the dynamic range of neuronal responses, and stimulus-specific clustering of population responses. The manner in which exposure enhanced the segregation of population responses into low-dimensional, stimulus-specific clusters suggested two main effects. First, stimulus responses became less variable across trials and more stereotyped, shrinking the radius of individual clusters. Second, responses to different stimuli became more distinct, increasing the distance between clusters. Using decoders we found that the information about stimulus identity was temporally specific, i.e. different time bins in the trial differed in their mapping, and visual exposure improved both variant and invariant aspects of stimulus encoding. Interestingly, we also observed a reduction in co-variability, similar to what has been previously documented in the context of attention and perceptual learning (27, 33, 34). In our data, the effect of exposure on spike-count correlations was complex: the strength of correlations was reduced with exposure, but knowledge about their structure remained beneficial for stimulus discrimination.

The brief stimulus presentations employed here resulted in a stereotyped, biphasic neuronal response in primary visual cortex, consisting of a high amplitude transient followed by a delayed persistent response. Stimulus decoding performance diverged from the expected dependence on firing rate, with accuracy peaking not on the response transient, but 200-400 ms after stimulus offset. Sustained and information-rich sensory responses, persisting beyond the period of sensory stimulation have been reported previously, not only under anesthesia, but in various sensory modalities and species in awake behaving animals. In the primary auditory cortex of awake marmosets, preceding stimuli suppressed or facilitated responses to succeeding stimuli for more than one second (35). In awake mice, the early sensory responses to a single brief whisker deflection encoded stimulus information, while the later activity appeared to drive the subjective detection (36). Notably,in awake mice primary visual cortex, an oriented flashing light induced a biphasic membrane voltage response that consisted of an early, transient depolarization and a delayed, slow depolarization (20). The delayed activity exhibited high orientation selectivity and influenced the evoked response to subsequent inputs in an orientation-selective manner. In awake macaques, a simultaneous change in both stimulus and background gave rise to a delayed V1 response that varied with the size of the background and correlated with the perception of a visual aftereffect (21). In human electroencephalography, information about a previously presented visual stimulus persisted even in the absence of delayed activity (activity-silent states) and could be decoded from an impulse response, long after stimulus presentation (37).

The persistence of stimulus information observed in our data and supported by the studies mentioned above, highlights a propensity for the primary visual cortex to maintain sensory information, far beyond the temporal intervals required by the traditional feed-forward model of the ventral stream. Instead, these findings are compatible with a dynamic coding framework for recurrent computation (29, 30, 38). In this framework, the cortical response to a stimulus emerges from an interaction between the input signals and the internal dynamical state of the network, including the ongoing activity (active states), but also the time-dependent properties of neurons and synapses (hidden states). Efficient recurrent processing relies on two simple requirements: (i) stimulus responses must persist beyond the duration of the stimulus, establishing a brief memory of recent events (fading-memory property) and (ii) the temporal evolution of network states in response to different stimuli must result in reproducible stimulus-specific trajectories (separability property). Both the memory and separability properties exhibited by a recurrent circuit can be optimized through plasticity by altering the network’s stimulus-response mapping (38).

The self-organized recurrent network model used here, builds on previous computational work showing that unsupervised changes in hidden states via local experience-dependent plasticity rules can increase performance on memory and prediction tasks (39) while matching numerous experimental findings on cortical variability (40). We employed response facilitation to boost the network’s response to brief stimuli during learning and considered its effect on network dynamics in several different implementations. We found that shifts in excitatory-inhibitory synaptic gains led to strong persistent responses after stimulus offset and over the exposure interval this boost in activity allowed for an increase in stimulus decoding performance. In contrast we found that while an increase in excitatory drive or shifts in neuronal excitability via intrinsic plasticity also led to strong persistent responses after stimulus offset, this boost in activity did not reliably improve stimulus decoding performance, suggesting that stable intrinsic network interactions are essential during learning. Notably, various other computational models have shown that the dynamics and performance of recurrent neural networks can be optimized via brain-inspired plasticity mechanisms. For example, spike-frequency adaptation was shown to expand the memory exhibited by recurrent circuits (41) and different forms of biologically plausible synaptic learning rules have been employed to enhance computational performance of recurrent networks in an unsupervised fashion (42–45). Furthermore, several studies made direct attempts to link learning in recurrent networks to optimization of state space dynamics (46, 47) or meta-learning (48). While neither of these recurrent models tried to explain how persistent responses after brief stimulation can interact with learning, they provide valuable insights into the various means by which refinements in internal network dynamics result in improved output performance.

The precise anatomical connectivity responsible for the observed reverberation of visual responses and the functional changes underlying exposure-driven improvements in stimulus discrimination are still unknown. Given the presence of both strong feedforward as well as, extensive feedback thalamo-cortical interactions (49, 50), we cannot exclude the possibility that exposure-driven changes in primary cortex responses originate from interactions with subcortical structures. In fact, studies have shown that slowly decaying inhibitory postsynaptic potentials in the lateral geniculate nucleus can maintain stimulus specific information for up to 300 ms and can modulate subsequent responses to reoccurring contours (51, 52). However, these effects were short lasting, while in our data repeated exposure to stimuli resulted in an increase in stimulus encoding across numerous trials and was associated with a strengthening of activation patterns in postexposure spontaneous activity. Interestingly, the activation of NMDA receptors in cortical layer 4, which receives the densest thalamocortical input, does not appear to be necessary for stimulus-selective response potentiation in V1 (53). An alternative is that exposure-dependent changes primarily affect local recurrent interactions within primary cortex and/or the long-distance recurrent interactions with higher cortical areas.

Vision depends on integrating the current sensory input in light of previous experience (54). This integration is achieved through the rich recurrent dynamics of the early visual system, which arise on the backbone of structural connectivity (55, 56). The connectivity of sensory areas is believed to capture the statistics of the environment, a process which would improve processing for expected stimuli (4, 7, 54, 57–59). However, exposure-driven changes in the primary sensory cortex of adult subjects, suggest a complex, occasionally divergent pattern of results. For example, adaptation classically leads to a reduction in response amplitude (10), and familiar stimuli can evoke reduced responses compared to novel stimuli (60). However, repetitive exposure can also alter the receptive fields and tuning of neurons in V1 (61), imprint responses to recent stimulus trajectories (24), increase the magnitude of responses to familiar sequences and signal predictions to missing elements (23). These intricate changes in response amplitudes with visual exposure appear to depend on many factors, among which, the frequency and duration of stimulation, the structure and complexity of the image set, the state of the animal and the precise signal measured. Regardless, the core findings outlined here do not rely heavily on changes in firing rates to familiar stimuli, but rather on the prolonged maintenance of stimulus information and the increase in discriminability with visual experience.

We interpret these results as an accumulation of evidence that optimizes the encoding of a stimulus set akin to learning a new stimulus statistics.

State changes are known to vary under anesthesia and can change the cortical response to stimulation. For example, deep anesthesia gives rise to alternating ‘up’ and ‘down’ states with distinct dynamic profiles that can have an impact on sensory coding. We did not observe such strong variations in our recordings, which were performed in a modified anesthetic protocol to mimic ‘awake’ like brain dynamics. Although a milder form of such variation occurs also during wakefulness, we found no systematic change in brain state that could trivially explain our results. Likewise, brief stimuli are known to generate robust responses irrespective of cortical state (62) which may explain the stability of sensory responses and brain states across recording sessions. Given that the experiments were performed under anesthesia, the reported exposure-driven changes in activity must involve “automatic” mechanisms, independent of attention and conscious control. Further work is necessary to determine to which extent these effects generalize to the waking state, where higher cortical areas with reciprocal connections to V1 as well as subcortical regions, such as the superior colliculus, thalamus and cerebellum are likely to play an important role in shaping V1 plasticity. In particular, top-down enhancement of task-relevant stimulus features and suppression of irrelevant ones, the level of attention, motivation and reward expectation, are all likely to guide learning-induced changes in V1.

Our study provides compelling evidence that repetitive visual exposure optimizes sensory processing in primary visual cortex, resulting in a better readout of stimulus-specific information. These findings suggest that the reliable visual discrimination of familiar stimuli can be partially achieved through separation of neuronal representations at the earliest cortical stage in the visual hierarchy. Future work should establish how these changes impact the transformation of sensory signals in the visual hierarchy, manifest at higher visual areas, and interact with behavioral states, such as attention or perception.

## Materials and Methods

### Electrophysiological recordings and data processing

Data was recorded from five adult cats (*felis catus*; mean age 2.7 years; range 1-5 years; two females) under general anesthesia during terminal experiments in two separate laboratories. The cats were bread internally, were housed together with other cats in small groups and experienced normal vision during development.All procedures complied with the German law for the protection of animals and were approved by the regional authority (Regierungspräsidium Darmstadt). For one of the cats, anesthesia was induced by intramuscular injection of Ketamine (10 mg/kg) and Xylazine (2 mg/kg) followed by ventilation with N2O:O2 (70/30%) and halothane (0.5%–1.0%). After verifying the depth of narcosis, pancuronium bromide (0.15 mg/ kg) was added for paralysis. Stimuli were presented binocularly on a 21-inch computer screen (HITACHI CM813ET) with 100 Hz refresh rate. To obtain binocular fusion, the optical axes of the two eyes were first determined by mapping the borders of the respective receptive fields and then aligned on the computer screen with adjustable prisms placed in front of one eye. Data was recorded with multiple 16-channel silicon probes from the Center for Neural Communication Technology at the University of Michigan (each probe consisted of 4 shanks, 3 mm long, 200 μm distance, 4 contact points each, 1,250 μm2 area, 0.3 — *0.5M*Ω impedance at 1 kHz). To extract multi-unit activity, signals were amplified 10,000 and filtered between 500 and 3500 Hz.

For four of the cats, anesthesia was induced by intramuscular injection of Ketamine (10 mg/kg) and Medetomidine (0.02 mg/kg) followed by ventilation with N2O:O2 (60/40%) and isoflurane (0.6%-1.0%). After verifying narcosis, Vecuronium (0.25mg/kg/h i.v.) was added for paralysis. Data was collected via multiple 32-contact probes (100 μm inter-site spacing, ≈ 1*M*Ω at 1 kHz; NeuroNexus or ATLAS Neuroengineering) and amplified (Tucker Davis Technologies, FL). Signals were filtered with a passband of 700 to 7000 Hz and a threshold was set to retain multi-unit activity. Thresholds remained fixed during data collection.

### Visual stimuli

Stimuli consisted of 34 shapes: 26 letters (A–Z) and 8 digits (0–7). They were white on black background and spanned approximately 5–7 degrees of visual angle.

More than 1700 trials (50 trails per stimulus) were recorded in every session. More than 6800 trials (200 trials per stimulus) were recorded in one of the sessions to test whether longer exposure leads to further improvements in stimulus encoding (Supp. Fig. 6).

Stimuli were presented in random order, i.e. consecutive trials corresponded to different stimulus conditions.

### Data analysis

Data was processed and analysed using custom code written in MATLAB (MathWorks). The Fieldtrip toolbox (63) was used for laminar analysis (Supp. Fig. 3).

### Spike-sorting

Spike sorting of the recorded multi-units was performed offline via custom software that computed principal components of spike waveforms in order to reduce dimensionality and grouped the resulting data using a densitybased clustering algorithm (DBSCAN). Only the well isolated clusters were considered single units and labeled separately. Spike-sorting resulted in 112 single units and 221 remaining multi-units, in total 443 units across all datasets.

### Current source density analysis

In 3 cats (7 sessions, 289 units), the recordings were performed with 32-channel linear arrays (100 micron spacing). Local field potentials (LFPs) to moving grating stimuli presented at maximal contrast were recorded either immediately before or immediately after the sessions with letters and digits. These LFPs were subject to current source density (CSD) analysis using a standard algorithm (64) based on the second spatial derivative estimate of the laminar local field potential time series. This analysis revealed successfully the short-latency current sink in the middle layers for each session, which has been shown to correspond most closely to layer 4 (65).

### Neuronal response properties

For every unit, the discriminability index *d*’, also known as Cohen,s effect size (25), for *n* stimuli, was calculated as:

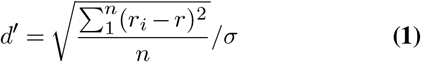

where *r_i_* is the mean response across trials to stimulus *i*, *r* is the mean response across all trials and *σ* is the common within population standard deviation, here 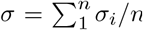, where *σ_i_* is the standard deviation of responses across trials to stimulus *i*. Note that low single-unit *d*’ values across many stimulus conditions (*n*=34) are expected, see for example (66) for reference.

The Fano factor was computed per unit according to Fano (67)

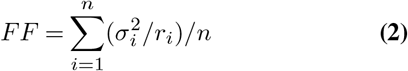

where *σ*, *r_i_* and *n* are defined as above.

Response sparseness *R*, describes the response distribution of a population of neurons to a single stimulus. Within each session, the response sparseness of the recorded population of neurons to each stimulus was calculated as:

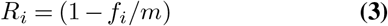

where *f_i_*/*m* is the fraction of units out of the total *m* that fired above the baseline in response to stimulus *i*.

Stimulus selectivity quantifies the responsiveness of a neuron across a set of stimuli and was defined as in (68). For each unit:

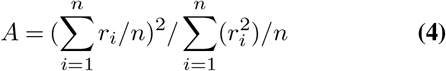

where *r_i_* is the unit response to stimulus *i* and *n* is the total number of stimuli. We used this measure in its inverted form *S*:

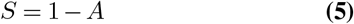

so that large values of S indicate high selectivity.

The response range *G*, was defined, for each unit, as the difference between the maximum and minimum response across all stimuli:

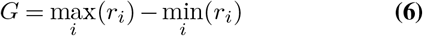

All of the measures defined above, were calculated for every 50 ms time-interval within the trial. When the reported values refer to larger time-windows, they represent averages over several 50 ms intervals.

### Stimulus classification

An instantaneous Naïve Bayes decoder was trained and tested on individual time bins of population responses. The size of a bin was 50 ms, unless specified otherwise. We performed cross-validation by randomly subsampling the data (k-1 data partitions used for training, 1 used for test, k repetitions; k=100). The task of the decoder was to determine the stimulus identity for each test trial, based on the population response in a particular time bin. Chance level was 1/number of stimuli = 1/34.

Support Vector Machines (SVMs) with quadratic kernels were applied using a similar cross-validation procedure for the computations shown in Supp. Fig. 7. For each data split, we trained the SVMs on either intact or trial-shuffled data (shuffling across trials within stimulus condition) and tested them on intact data to test whether access to the correlation structure present in the data leads to better performance.

### Self-organizing recurrent network (SORN)

The neural network model was composed of 80% excitatory (*N^E^* = 200) and 20% inhibitory units (*N^I^* = 50). Connectivity matrices *W^IE^*, *W^EI^* and *W^II^* were dense, randomly drawn from the interval [0,1] and normalized so that the incoming connections to each neuron sumed up to a constant 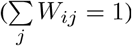.

The connections between excitatory units *W^EE^* were random and sparse and followed soft topological constrains (*p^EE^* = 0.1 was the connection probability for neighboring units, i.e. every 10 consecutive units were considered neighbors; *p^EE^* = 0.01 was the connection probability for non-neighbors). The threshold values for excitatory (*T^E^*) and inhibitory units (*T^I^*) were drawn from a uniform distribution in the interval [0, 0.5] and [0, 0.3]. The network state at time t was given by two binary vectors *x*(*t*) ∈ 0, 1*^N^E*, and *y*(*t*) ∈ 0, 1*^N^I*, representing activity of the excitatory and inhibitory units, respectively. Each timestep *t* corresponded to ≈ 20*ms* of real time.

The network evolved using the following update functions:

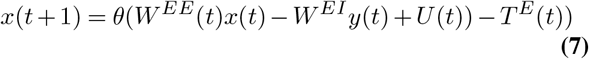

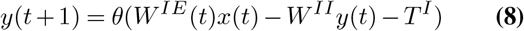

The Heaviside step function *θ* constrained the network activation at time *t* to a binary representation: a neuron fired if the total drive it received was greater than its threshold.

The stimulus set was composed of 10 digits, 6×6 pixels each. Every 5th excitatory unit received input from one corresponding image pixel, i.e. 36 units were input units, 164 units were reservoir units. The input *U*(*t*) varied as a function of time (blue marking in Figure 6): initially it represented the luminance of the stimulus at particular pixel location (“on” response, 2 time steps), later it represented half the luminance of the reversed stimulus image at the same location (“off” response, 7 time steps). For learning, we utilized a simple additive spike-timing dependent plasticity rule that increased (or decreases) the synaptic weight *W^EE^* by a fixed amount *η*_STDP_ = 0.001 whenever unit *i* is active in the time step following (or preceding) activation of unit *j*.

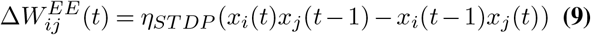

In addition, synaptic normalization was used to proportionally adjust the values of incoming connections to a neuron so that they summed up to a constant value *c_E_* = *c_I_* = 1.

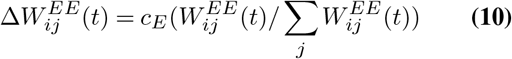

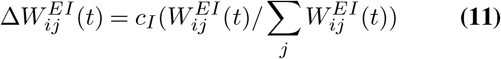

To stabilize learning, we used a homeostatic intrinsic plasticity (IP) rule that spread the activity evenly across units, by modulating their excitability using a learning rate *η_IP_* = 0.001. At each timestep, an active unit increased its threshold, while an inactive unit lowered its threshold by a small amount, such that on average each excitatory neuron fired with the target firing rate *μ_IP_* = 0.1:

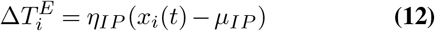

Response facilitation was applied for a fixed time interval in each trial (20 time steps, shaded area in Figure 6). Four different implementations were considered: (1) the incoming synaptic inhibition was lowered (*m_I_* = 0.5); (2) the incoming synaptic excitation was increased (*m_E_* = 1.5); (3) an additional noisy input was fed to all excitatory units (uniformly distributed in the interval [0, 0.01]); (4) the target firing rate of excitatory units set via intrinsic plasticity was increased (*η_IP_* = 0.2).

The SORN model was implemented in MATLAB. A similar implementation of the model in Python (with absent *W^II^* connections) can be found at https://github.com/chrhartm (40).

## AUTHOR CONTRIBUTIONS

Conceptualization, A.L., W.S. and D.N.; Methodology, A.L., and D.N.; Investigation, A.L., C.L., and D.N.; Writing – Original Draft, A.L. and D.N.; Writing – Review & Editing, A.L.; C.L. and W.S.; Funding Acquisition, P.F., W.S., and D.N.; Resources, P.F., W.S., and D.N.; Supervision, W.S., and D.N.

## ACKNOWLEDGEMENTS

Special thanks go to Thomas Wunderle and Jianguang Ni for technical assistance during recordings. DN and AL acknowledge grant support by the Deutsche Forschungs-Gemeinschaft (DFG NI708/5-1). PF acknowledges support by DFG (SPP 1665), EU (FP7-604102-HBP). WS acknowledges the Reinhart Kosselleck grant of the German Research Foundation and the EU’s 7th Framework Programme (FP7/2007-2013 Neuroseeker).

