## Supplementary Figures for "Visual exposure enhances stimulus encoding and persistence in primary cortex"

Summary

The supplementary material consists of the following figures:


[S1] Stability of cortical signals during recordings

[S2] Impact of exposure on firing amplitude and variability

[S3] Impact of exposure in different cortical laminae

[S4] Response clustering in low dimensional projections

[S5] Exposure-driven increase in stimulus classification

[S6] Classification increase with additional exposure time

[S7] Effects of visual exposure on spike-count correlations

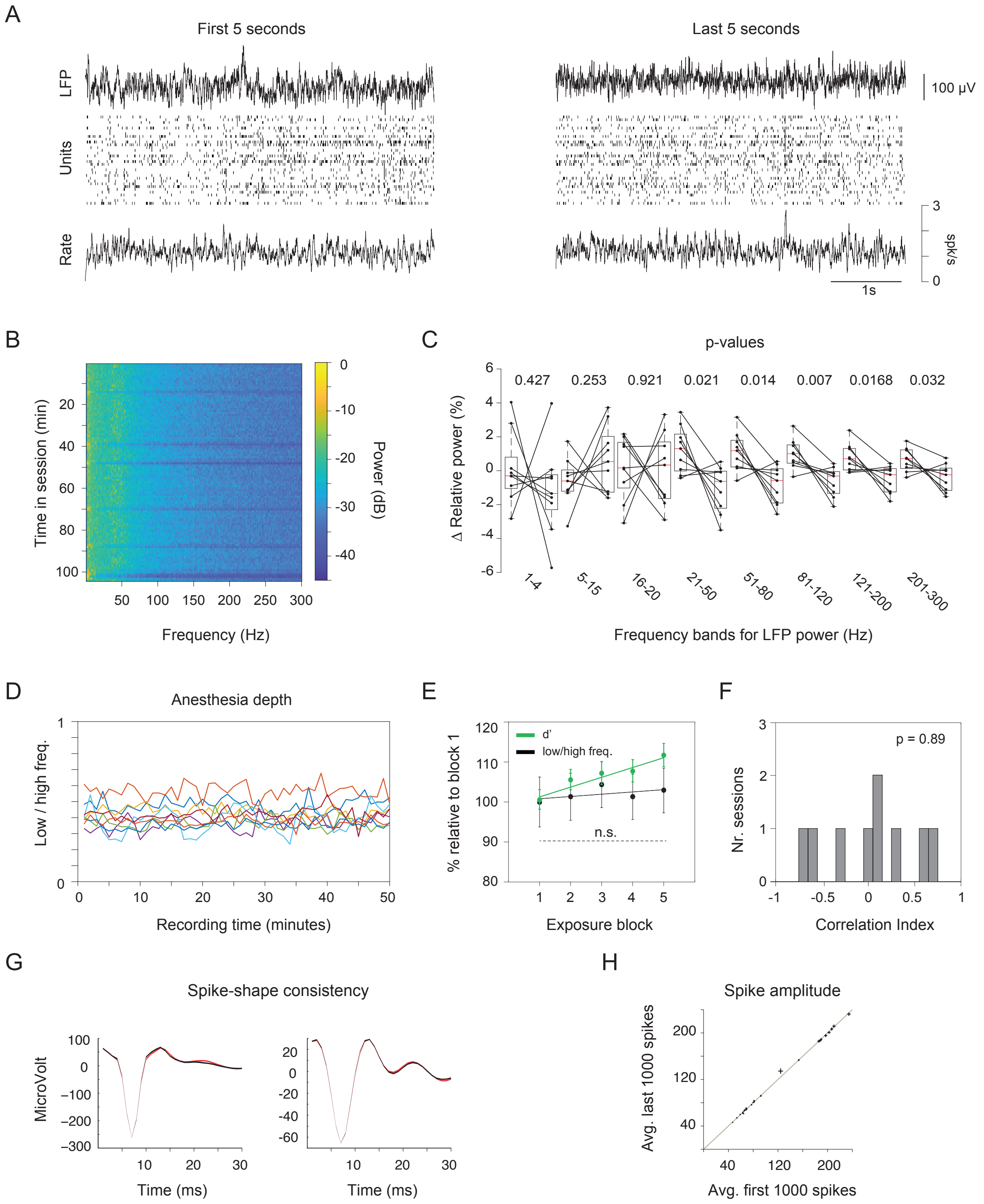


**Supp. Fig. 1** Stability of cortical signals during recordings: A) Example local field potential (LFP) traces and activity of simultaneously recorded units from the first and last five seconds in a recording session B) LFP power spectrum over the course of a session displays small fluctuations and no apparent trend. C) Relative change in LFP power in different frequency bands (9 sessions) is small (several percentages) and not significant (p-values for t-test between early and late power are indicated above boxplots). While a number of frequency bands show a trend towards change, none survives multiple-comparison correction and they are not consistent with a trivial change in state over the course of exposure (both low and high frequency power trends towards decrease over exposure). This suggests that the cortical state, while variable, does not change systematically across the recordings. D) Ratio of LFP power (low 1-5Hz/ high 20-300 Hz) in 9 sessions shows no systematic changes with exposure. E) d' changes gradually with exposure in the session, but the ratio of LFP power does not (d' linear fit $y = 2.4x + 98.66$ in green, linear trend $p = 0.009$; LFP power ratio linear fit $y = 0.57x + 100.24$ in black, linear trend $p = 0.34$ n.s.). F) Changes in LFP ratio do not show structured correlation with changes in d' over five exposure blocks across individual sessions. G) Average spike shape of first 1000 spikes (black) vs. last last 1000 spikes (red) for two example units. Shaded area indicates s.e.m. H) Spike amplitude of first vs. last 1000 spikes for all units in a session. Cross lines indicate s.e.m.

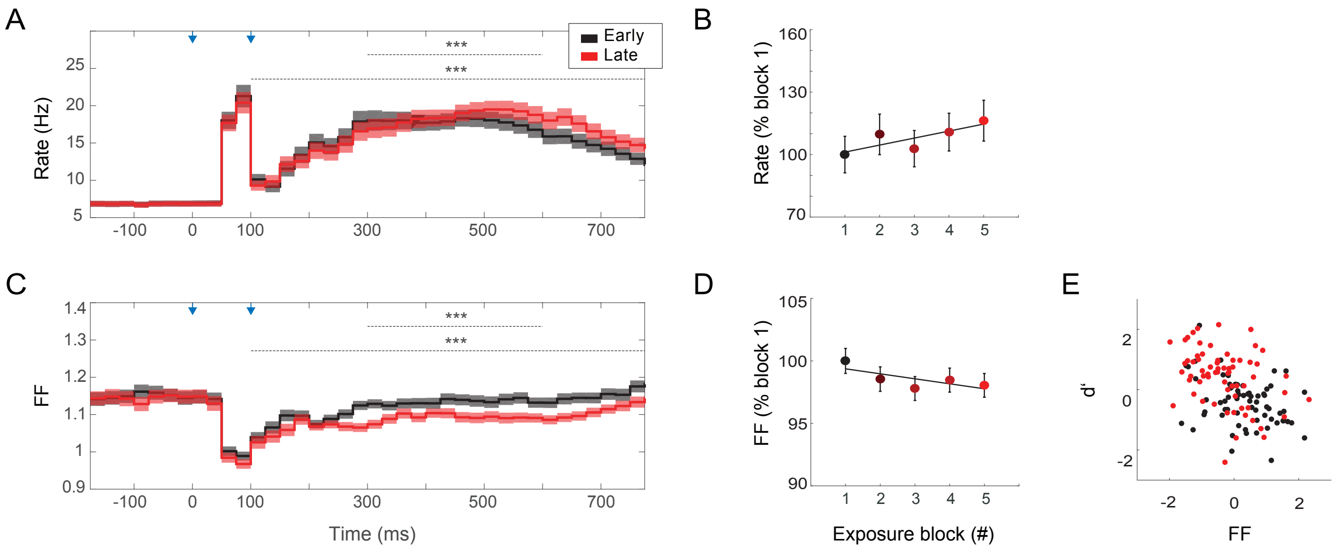


**Supp. Fig. 2** Impact of visual exposure on firing rate amplitude and variability. A) Mean firing rate amplitude across 443 recorded units over the course of the trial increases significantly with visual exposure (margins depict s.e.m.; paired t-test across 443 units shows significant differences in rate; p<0.001 for the intervals 100-800 ms and 300-600 ms). B) Firing rates increase over five exposure blocks (error bars depict s.e.m.). Differences between block 1 and 5 are significant but linear trend is not (16.28% increase between block 1 and 5, paired t-test p = 4.1e-05, t = -4.14, df = 442; linear fit, y = 3.36x + 97.8, linear trend p = 0.09). C) Mean firing variability (Fano factor) across 443 recorded units over the course of the trial decreases significantly with visual exposure (paired t-test, p < 0.001 for the intervals 100-800 ms and 300-600 ms). D) Firing variability decreases over five exposure blocks. Differences between block 1 and 5 are significant but linear trend is not (2% decrease between block 1 and 5, paired t-test p = 0.01, t = 2.44, df = 442; linear fit, y = -0.39x + 99.7, linear trend p = 0.1). E) Normalized average firing variability across the neuronal population (interval 300-600 ms, six consecutive 50 ms temporal bins) against the corresponding discriminability scores (d'), superimposed for all recording sessions. Similar variability levels resulted in higher d' scores in late trials (red) compared to early trials (black) in a session.

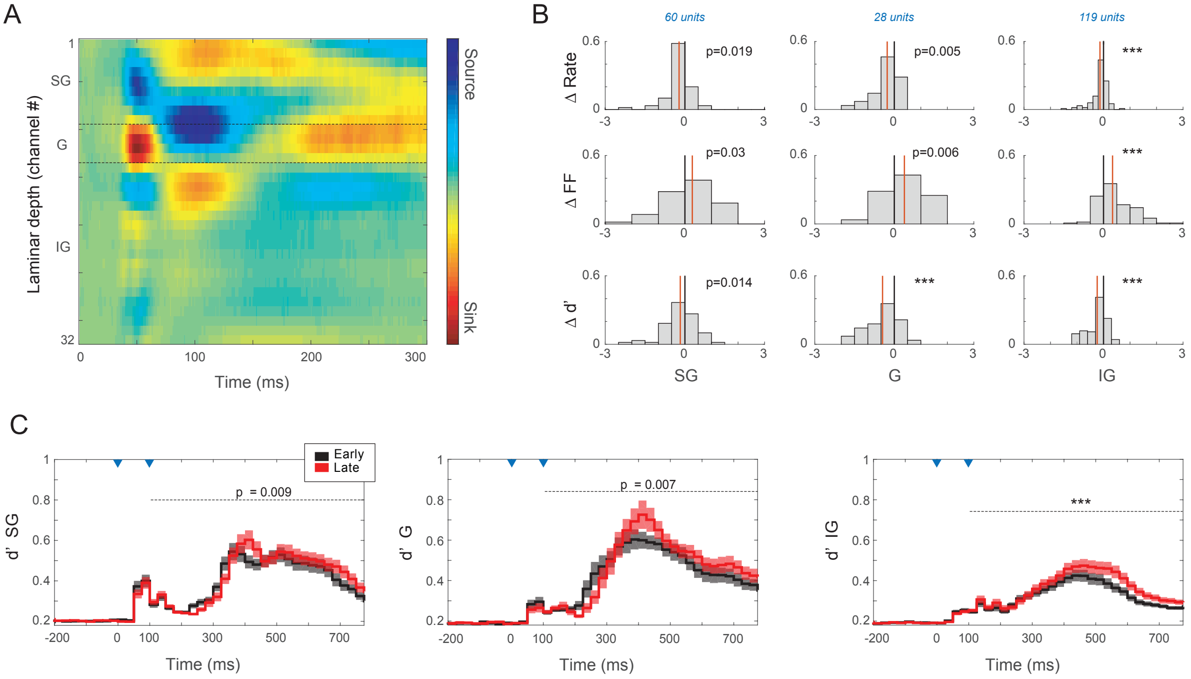


**Supp. Fig. 3** Impact of visual exposure on unit responses from different cortical laminae. A) Current source density derived from a 32-contact laminar array allows the determination of laminar structure (example session). B) Histograms of differences in firing rate, firing variability and $d'$ between early and late trials, separately for units from different cortical laminae (data from 7 sessions recorded with 32-channel laminar arrays, 207 multiunits; values were z-scored per session; paired t-test for the 300-600 ms time-interval). C) Stimulus discriminability over the course of the trial in different cortical laminae. Significance shown for 100-800 ms. *** stands for p<0.001.

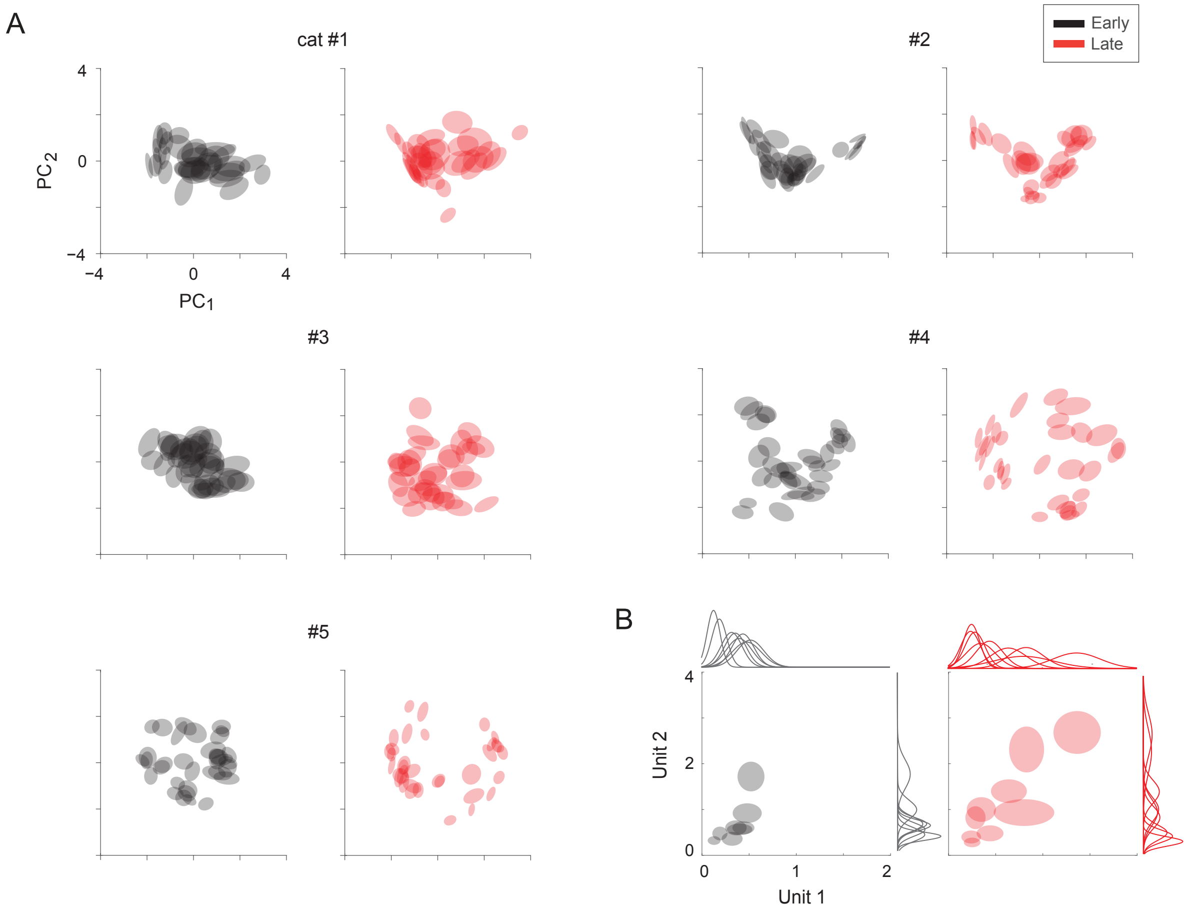


**Supp. Fig. 4** Visual exposure increases the stimulus-specific clustering of population responses in low dimensional projections. Example sessions from five recorded animals. Principle component analysis was performed separately for early (black) and late (red) trials in each session. The centers of the depicted ellipses correspond to the mean population responses to 34 visual stimuli in the projection space described by the first two principle components. Each ellipse circumscribes the data points within one standard deviation from the mean. For all sessions, the PCA was applied for spike-count population vectors over a fixed 50 ms time bin, 400 ms after stimulus offset. B) Impact of exposure can be seen for pairs of multi-units with overlapping stimulus-response preferences, even without projection in PC space (spike-counts over 50 ms). In this example, both units increase their response range with exposure and exhibit an increased segregation of stimulus responses. As above, the ellipses circumscribe the data points within one standard deviation from the mean.

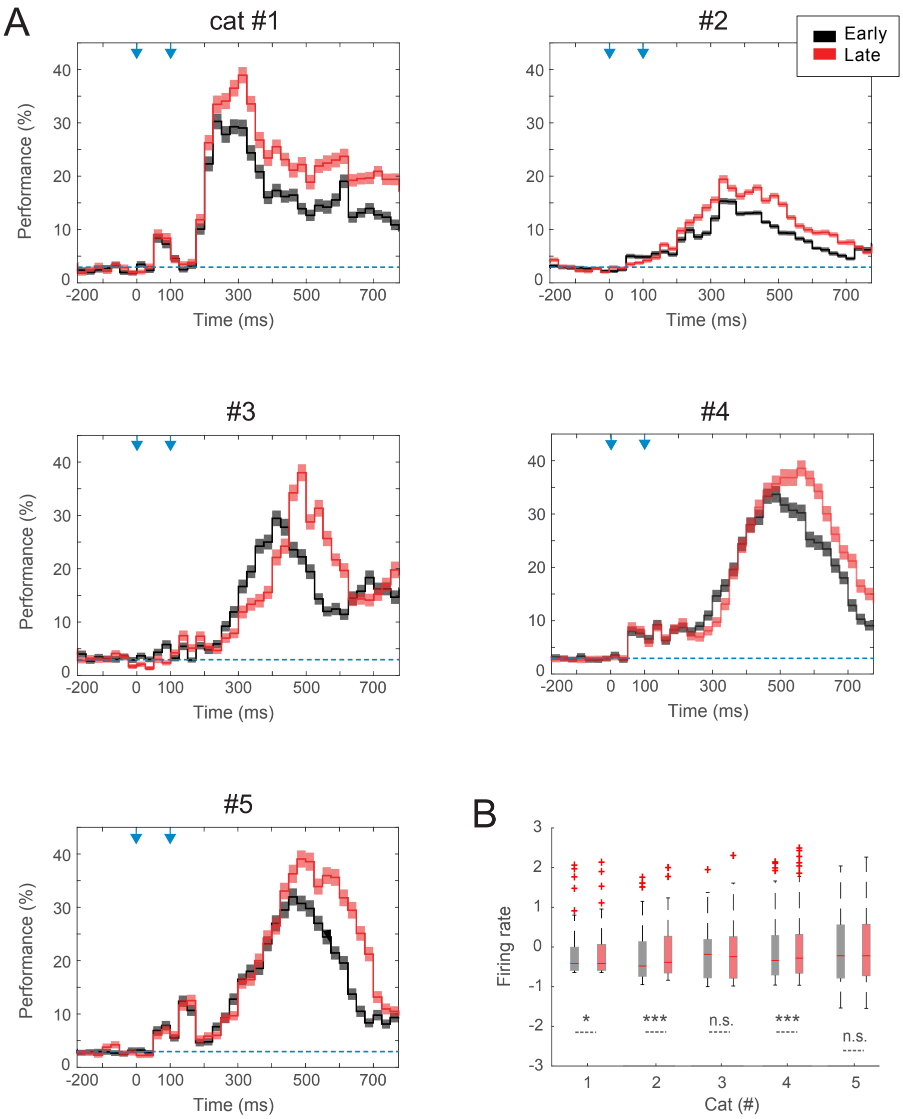


**Supp. Fig. 5** Visual exposure increases stimulus classification performance (5 cats; 50 ms bins, 25 ms sliding bin; 34 stimuli, chance level 2.94 %). Average performance of a Naïve Bayes classifier was higher for late trials (red) compared to early trials (black) in every animal (increase in peak-performance 13-59.6% increase, two-tailed t-test, all p-values<0.001; increase in performance-AUC 14-64%, 100-800ms interval, two-tailed t-test, all p-values<0.001) B) Comparatively, firing rate changes were modest (box-plots show z-scored values across units in separate animals; paired t-test * p<0.05; *** p<0.001). For cat 3 and 5 the firing rate changes with exposure were not significant.

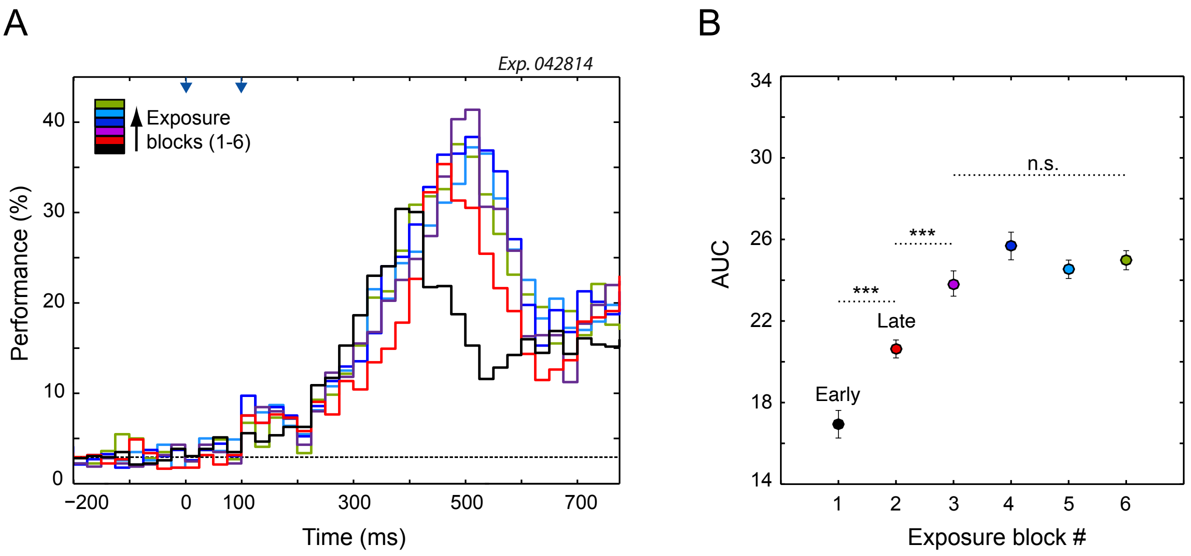


**Supp. Fig. 6** Classification performance increases with additional exposure time. A) Performance profile of instantaneous Bayesian classifier over the course of the trial for four additional blocks of exposure (25 stimulus repetitions per block, color indicates block order). B) Difference in performance-AUC between block 1 and block 2 was strong (21% increase, interval 300-600 ms; early trials black, late trials red). Longer exposure led to a further performance-AUC increase (15.5% increase between block 2 and block 3). Significance tests were applied for differences across trial-blocks (ANOVA with Bonferroni correction for multiple comparisons). Blocks 1 and 2 were significantly different from each other and all other blocks (p-values<0.001). Blocks 3,4,5 and 6 were significantly different from blocks 1 and 2 (p-values<0.001) but not each other suggesting that the performance may have saturated after $\approx$ 75 repetitions per stimulus.

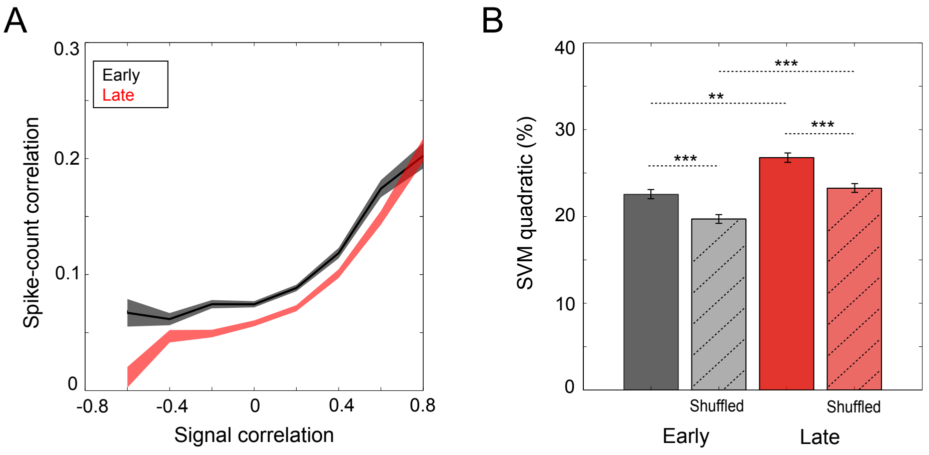


**Supp. Fig. 7** Effects of visual exposure on spike-count correlations (SCCs. A) Mean SCCs across trials increase as a function of signal correlations, for both early (black) and late (red) trials (average across all simultaneously recorded pairs, 300-600 ms interval, margins indicate s.e.m.). SCCs are reduced by visual exposure (insert indicates shift in the distribution of spike-count correlations). B) Peak performance of a Support Vector Machine with quadratic kernels trained on original data (intact correlation structure, black and red) or shuffled data (compromised correlation structure, gray and yellow) and tested on original data (error bars indicate s.e.m.). Shuffling was performed across trials, separately within each stimulus condition (i.e. signal correlations were not affected). Both the effects of visual exposure and shuffling were significant (two-way ANOVA with Bonferroni correction, ** p<0.01, *** p<0.001).
